# Reduced B cell activation in Germinal center reaction of the mouse model of Li-Fraumeni syndrome

**DOI:** 10.64898/2026.01.21.700937

**Authors:** Koichi Hasegawa, Jacqueline H Barlow, Santosh K Gothwal

## Abstract

The *TP53* mutation associated with Li-Fraumeni syndrome in humans is known to exhibit p53 gain-of-function properties leading to early cancer onset. To explore whether these patients also have compromised immune responses due to p53 mutations, we investigated the humoral immune response using a Li-Fraumeni mouse model harboring a structural mutation, *Trp53R172H* (equivalent to codon 175 in humans). *Trp53R172H* mice were immunized with sheep red blood cells, and the Germinal center response was monitored. Our results revealed that SRBC-immunized *Trp53R172H* mice exhibit reduced B cell activation during the GC reaction. These suggest a selective role for p53 in promoting B cell activation early in the GC reaction prior to BCL6 upregulation. We propose that impaired B cell activation in Li-Fraumeni patients could contribute to immune deficiencies and heightened susceptibility to autoimmune disorders, potentially influencing the development of secondary cancers and impairing therapeutic responses to chemotherapies and immunotherapies.

**Key Points:**

1. *Trp52R172H* mice exhibit reduced B cell activation during the Germinal center reaction
2. Ratio of activated B cells in DZ to LZ remain unchanged in *Trp52R172H* mice.

## Introduction

A key step of infection clearance is naïve B cell presentation of antigens to T cells located in secondary lymphoid organs [1]. This interaction requires naive B cell activation leading to the B cell receptor (BCR) activation and BCL6 expression [2]. Germinal center (GC) B cells express activation induced cytidine deaminase (AID) which induces DNA-double strand breaks in GC B cells and activates the DNA damage response [3, 4]. BCL6 plays a necessary role to promote the survival of activated GC B cells in the dark zones (DZ) and the light zones (LZ) [5]. Specifically, BCL6 suppresses the DNA damage response, p53 regulation, and apoptotic signals during the GC reaction to promote cell survival {Basso, 2010, 20510734;Basso, 2010, 20510734.

The tumor suppressor p53 regulates the DNA damage response, proliferation and tumorigenesis [6, 7]. Indeed, p53 has been proposed to inhibit class switch recombination (CSR) in B cells [8]. However, whether p53 plays a role in GC B cell activation, a step prior to its suppression by BCL6, remain unknown [5, 9, 10]. A key transition point of p53 functions in the GC reaction would be at the beginning of GC reaction where BCL6 activity is minimal, thus p53 levels could impact B cell activation. Soon after the GC reaction ends and BCL6 levels sharply decline in GC B cells, p53 functions might be upregulated in GC B cells harboring a high affinity BCR and are committed to differentiation into memory and plasma B cells [9, 11]. Therefore, p53 may function in GC B cells at two steps, first during the B cell activation and second in GC B cells undergoing terminal differentiation .

Li-Fraumeni syndrome is a cancer predisposition syndrome caused by germline mutations in the *TP53* tumor suppressor gene [12]. Li-Fraumeni syndrome is characterized by a wide spectrum of cancers with early onset including soft-tissue sarcomas, osteosarcomas, adrenocortical carcinomas (ACC), central nervous system tumors, and very early onset female breast cancers occurring before 31 years [12]. Germline *TP53* variants associated with Li-Fraumeni Syndrome have been estimated to be present in 50–80% of children with ACC [13-15] or choroid plexus carcinomas [13, 16] and up to 73% of children with rhabdomyosarcoma of embryonal anaplastic subtype [17] and between 3.8 and 7.7% in females with breast carcinoma before age 31 [18]. Despite these clinical studies on Li-Fraumeni patients, it is still unclear whether the gain of p53 functions in these patients impacts their humoral immune response.

In this study, we use a mouse model of Li-Fraumeni syndrome to examine the role of the gain-of-function *Trp523R172H* mutation in GC B cells. Our results show that *Trp53R172H* heterozygous mice have defective B cell activation in response to sheep red blood cell (SRBC) immunization, suggesting a role for p53 in normal GC B cell activation. Our results highlight a critical role of p53 in assuring the proper GC reaction establishment and suggest a reduced humoral immune response in Li-Fraumeni patients leading to defective immune responses in these patients.

## Results

### Generation of *Trp53R172H* by allelic recombination

Mice harboring one copy of the *Trp53R172H* allele were treated with tamoxifen to induce the recombination of Lox-Stop-Lox (*rLSL*) sequence in the first intron on *Trp53* locus, which induces *Trp53R172H* expression (Figure 1A). The recombination status was assessed by PCR amplification of genomic DNA. Due to the similar sizes of PCR products from the two alleles (290 bp for WT vs 330 bp for *Trp53R172H* allele), recombination frequency was estimated by quantifying PCR band intensities. Of the 12 mice treated with tamoxifen, two showed no detectable PCR amplification, and 4 exhibited a ratio of *rLSL* to wild-type allele band intensity of less than 0.90. The remaining six mice, which showed an *rLSL*-to–wild-type band intensity ratio greater than 0.90, were included in subsequent analyses (Figure 1C). This indicates successful establishment of the heterozygous *Trp53R172H* mice (*Trp53R172H* mice hereafter) for the GC reaction analysis.

**Figure 1:**
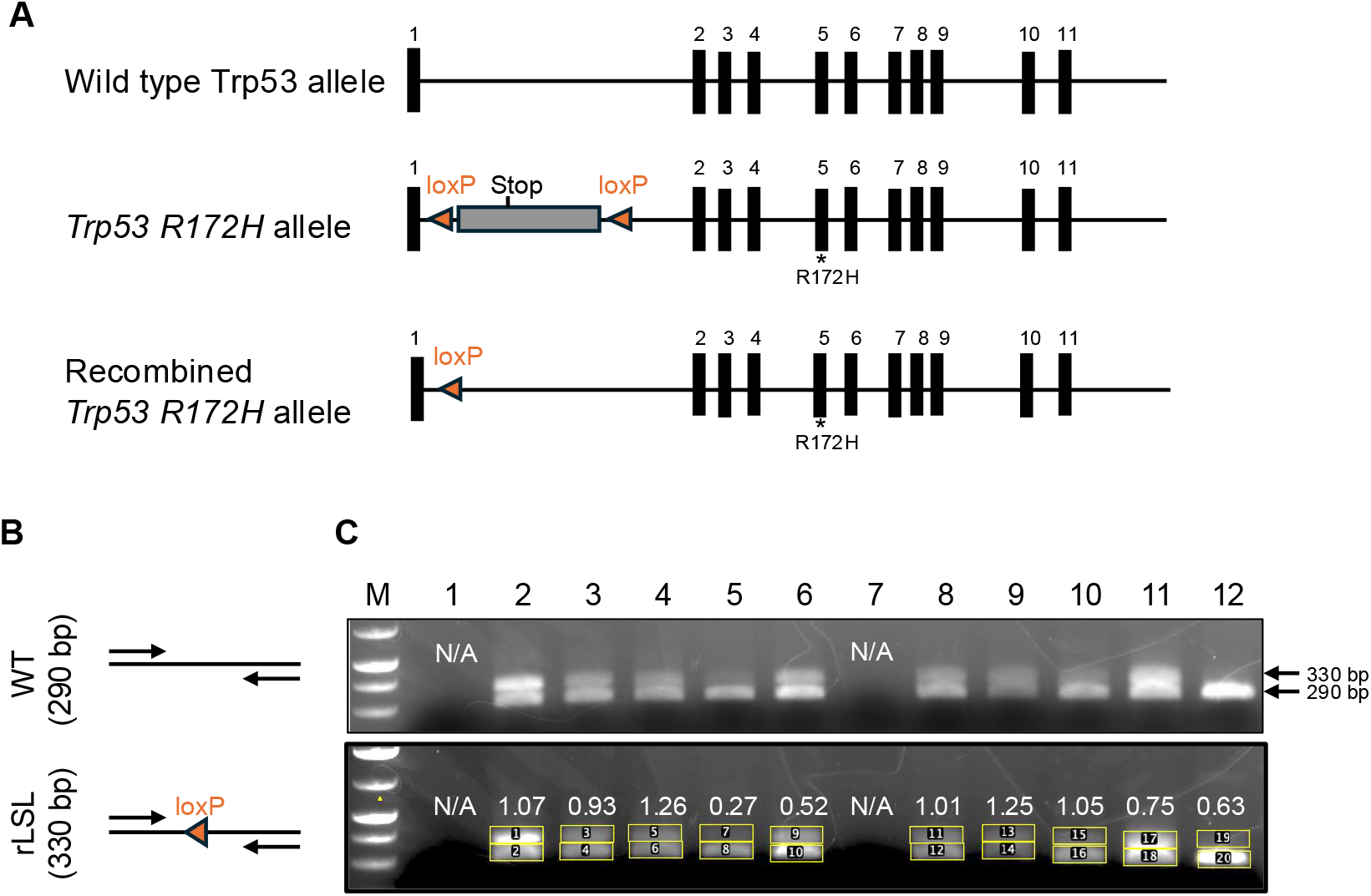
Allelic configuration and PCR genotyping of wild-type and conditional *Trp53 R172H* mice. **(A)** Schematic representation of the wild-type and mutant Trp53 (R172H) alleles. After Cre-loxP recombination, the R172H locus is transcribed. **(B)** Schematic representation of primer locations for detecting the wild type and the recombined LSL (*rLSL*) allele in the tamoxifen-induced *Trp53 R172H* mice. **(C)** Agarose gel showing PCR products of the control WT (290 bp) and *rLSL* (330 bp) alleles from *Trp53 R172H* mice.

### Reduced B cell activation in the *Trp53R172H* mice

To determine whether *Trp53R172H* mice exhibit an altered GC response, we analyzed SRBC-induced GC reactions in 11 control and six *Trp53R172H* mice (Figure 2A). We first quantified splenic B cell numbers by staining for the B cell marker B220. The average number of B220^+^ B cells was comparable between control and *Trp53R172H* mice (383,579 ± 3,603 vs. 383,248 ± 3,816, respectively; Figure 2B), indicating that *Trp53R172H* mice do not exhibit altered total B cell numbers following SRBC immunization.

**Figure 2:**
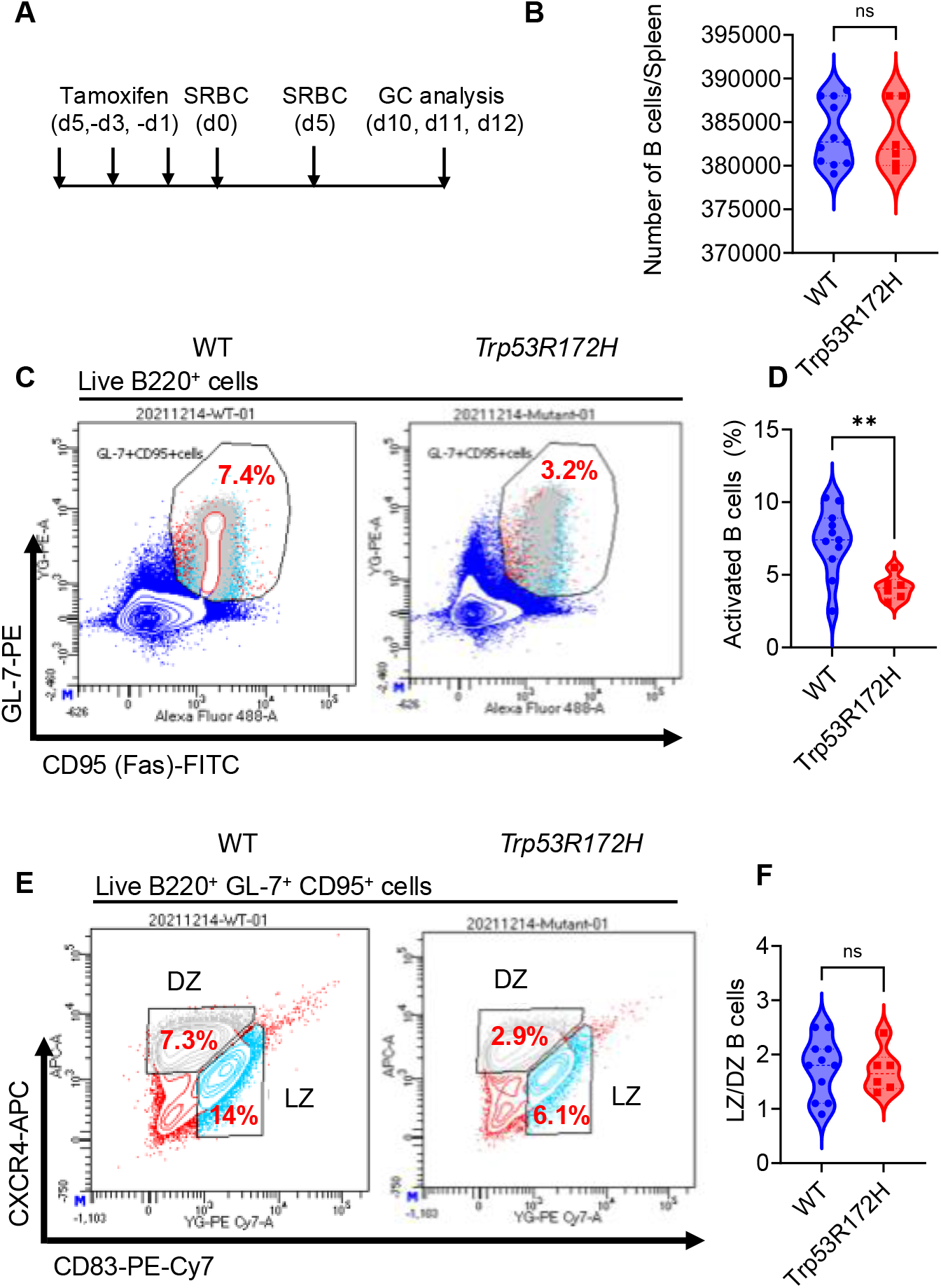
Reduced B cell activation in SRBC-induced GCs of *Trp53 R172H* mice. **(A)** Schematic of tamoxifen injection and SRBC-induced germinal center (GC) reaction in control and *Trp53 R172H* mice. **(B)** The total number of B cells in control (n=11) and *Trp53 R172H* mice (n=6) is shown. **(C)** Representative FACS plots identifying activated GC B cells in control and *Trp53R172H* mice. Among B220^+^ B cells, activated GC B cells were defined as GL7^+^ CD95^+^. **(D)** The proportion of activated B cells in control and *Trp53 R172H* mice as a fraction of total splenic B cells is shown. **(E)** Representative FACS plots identifying the dark zone (DZ) and light zone (LZ) population in control and *Trp53 R172H* mice. **(F)** Ratio of LZ B cells to DZ B cells in the control and *Trp53 R172H* mice is shown. (unpaired t-test). SRBC;Sheep red blood cells, GC; germinal center, DZ; dark zone, LZ; light zone.*p<0.05, **p<0.01

To assess B cell activation, we next gated on B220^+^ cells expressing the GC activation markers CD95 (Fas) and GL7. Notably, the fraction of activated GC B cells was reduced by approximately 50% in *Trp53R172H* mice compared with control mice (4.1% ± 0.8% vs 7.3% ± 2.3% respectively; Figure 2C, D), indicating impaired B cell activation in the *Trp53R172H* mice. Given the reduction in activated GC B cells, we next examined whether GC zoning dynamics were affected. We analyzed the distribution of GC B cells between the dark zone (DZ; CXCR4^high^ CD83^low^) and light zone (LZ; CXCR4^low^ CD83^high^). The LZ/DZ ratio was comparable between control and *Trp53R172H* mice (Figure 2E, F), indicating that cyclic migration between GC compartments is preserved despite reduced B cell activation in the *Trp53R172H* mice. Together, these data demonstrate that *Trp53R172H* mice exhibit impaired GC B cell activation without detectable disruption of GC zonal organization.

### Reduced accumulation of GC B cells in the DZ and LZ compartments of *Trp53R172H* mice occurs due to defective B cell activation

Activated GC B cells cycle between DZ and LZ compartments to undergo somatic hypermutation (SHM) and affinity maturation [19, 20]. Given the reduced frequency of activated GC B cells in *Trp53R172H* mice (Figure 2C, D), we further examined whether the representation of GC B cells in the DZ and LZ compartments was altered when normalized to total splenic B cells versus activated GC B cells. The proportion of DZ and LZ B cells relative to total splenic B cells was significantly reduced in *Trp53 R172H* mice (Figure 3A–B). However the frequency of DZ and LZ B cells relative to activated B cells was unchanged between control and *Trp53R172H* mice (Figure 3C–D). Together, these results indicate that the reduced representation of DZ and LZ B cells in *Trp53R172H* mice is primarily attributable to decreased B cell activation rather than impaired entry or maintenance of activated B cells within the DZ and LZ compartments.

**Figure 3:**
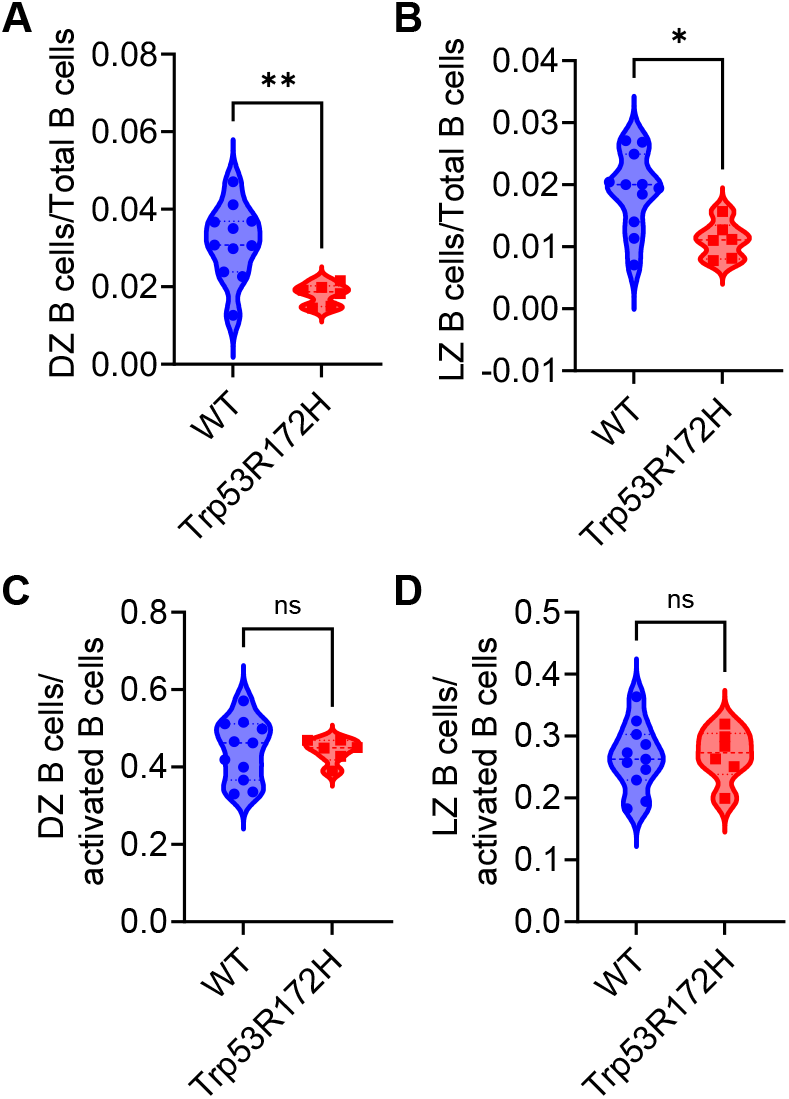
Reduced accumulation of GC B cells in the DZ and LZ compartments of *Trp53R172H* mice occurs due to defective B cell activation. **(A)** The proportion of DZ B cells relative to the total splenic B cells in control and *Trp53 R172H* mice is shown. **(B)** The proportion of LZ B cells relative to the total splenic B cells in control and *Trp53 R172H* mice is shown. **(C)** The proportion of DZ B cells relative to the activated splenic B cells in control and *Trp53 R172H* mice is shown. **(D)** The proportion of LZ B cells relative to the activated splenic B cells in control and *Trp53 R172H* mice is shown (unpaired t-test). DZ; dark zone, LZ; light zone. *p<0.05, **p<0.01.

## Discussion

In this study, we analyzed the GC reaction in control and *Trp53 R172H* gain-of-function mutant mice. The total number of splenic B cells was similar between genotypes (Figure 2B); however, *Trp53R172H* mice exhibited reduced B-cell activation (Figure 2A–D), indicating that p53 promotes the GC reaction by enhancing B-cell activation. This indicates that gain of p53 functions in Li-Fraumeni patients may impair proper humoral immune response due to defective GC assembly.

There are several key steps to activate GC B cells. Firstly, activation defect may reflect impaired regulation of genes involved in B-cell receptor (BCR) and T-cell signaling, which support T-cell cytokine secretion that enhances BCR-mediated B cell activation in the GCs [21, 22] (Figure 4A). Alternatively, defective macrophage-mediated antigen delivery to the subcapsular sinus of secondary lymphoid organs may compromise antigen presentation to GC B cells in *Trp53R172H* mice and in Li-Fraumeni patients [23, 24], weakening T–B interactions and B-cell activation [25]. It has been shown that *CCR7* is a direct transcriptional target of p53 [26], and is highly expressed in GC B cells. Therefore, the reduced B cell activation in *Trp53R172H* mice may result from impaired CCR7-mediated migration of B cells toward the T–B cell border where they present antigens to cognate T cells, a necessary step for B cell activation [24, 27]. Future studies comparing gain-of function mutations to knockout p53 alleles will help determine whether reduced B cell activation is due to expression of the *Trp53R172H* p53 variant or loss of p53 heterozygosity.

**Figure 4.**
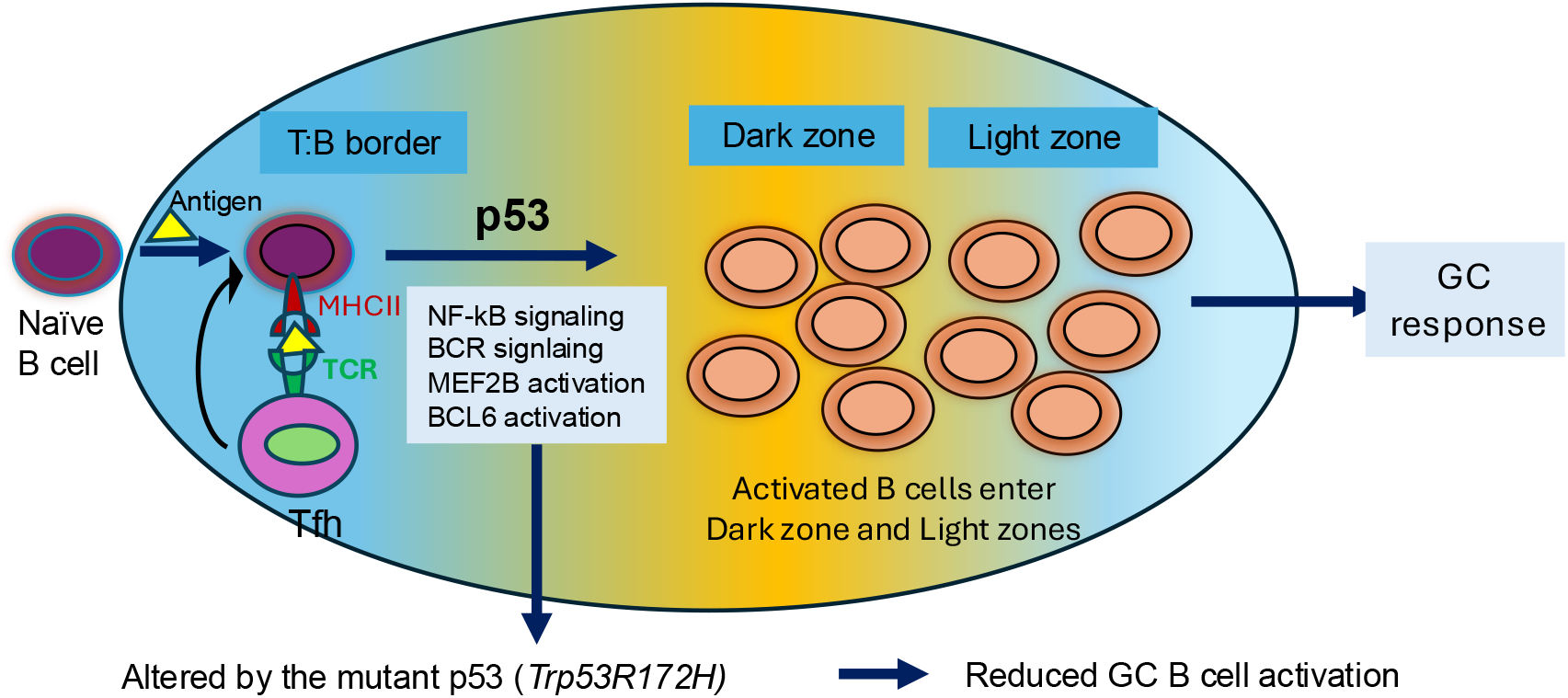
Altered regulation of p53 on genes involved in B cell activation during the GC reaction. Schematic representation of T and B cell interactions within the marginal zones of secondary lymphoid organs, occurring prior to the initiation of the GC reaction and BCL6 expression. The p53-dependent signaling in both T and B cells could contribute to proper B cell activation via several signaling mechanisms (NF-κB, BCR signaling), ultimately resulting in optimum GC response. Deficiency of wild type p53 function in *Trp53R172H* mice leads to reduced B cell activation that could impair the GC response and impair the humoral immune response.

Our observations of reduced B cell activation in *Trp53R172H mice* suggest that p53 may regulate B-cell activation prior to BCL6 expression, as BCL6 directly suppresses p53 [28]. Therefore, p53 function in B cell activation could occur in parallel with MEF2B activation, at an early stage of GC B cell activation.[29]. It might be possible that p53 loss of heterozygosity or gain of function could indirectly affect MEF2B regulation, resulting in reduced B cell activation [29-32]. Given that MEF2B activity is epigenetically regulated by histone deacetylases (HDACs) [33] and p53 antagonizes HDAC function [34], altered chromatin accessibility at MEF2B target loci in *Trp53R172H* mice may contribute to reduced B-cell activation. Consistent with this, gain-of-function p53 mutations associated with Li-Fraumeni syndrome have been shown to transcriptionally regulate MEF2 family target genes[35]. Future studies are needed to determine whether p53 gain-of-function directly modulates MEF2B-driven transcriptional programs during B-cell activation.

In summary, our findings suggest that humoral immunity may be compromised in patients with Li-Fraumeni syndrome due to gain of p53 functions and subsequently impaired B-cell activation [36, 37], potentially limiting the production of high affinity antibodies in these patients [38]. Our results highlight a multifaceted role for p53 in B-cell activation and the GC reaction, with its dysregulation contributing to immune defects in Li-Fraumeni syndrome.

## Author contribution

KH performed experiments, analyzed the data and wrote the manuscript. JHB critically reviewed the manuscript, provided constructive feedback, and wrote the manuscript. SKG designed the project, received funding, performed experiments, analyzed the data and wrote the manuscript.

## Acknowledgment

This work was supported by the Japan Society for the Promotion of Science (JSPS) under grant 21K16142 awarded to SKG.

## Material and Methods

### Mice

B6.129-Gt(ROSA)26Sortm1(cre/ERT2)Tyj/J (Jackson Laboratories, 008463) mice were crossed with B6.129S4(Cg)-Trp53tm2.1Tyj/J (Jackson Laboratories, 008183) mice to generate double heterozygous R26-CreER^T2^/*Trp53*^*R172H*^ mice, hereafter referred to as *Trp53 R172H* mice. Tamoxifen-untreated *Trp53 R172H* mice were used as controls in this study.

### Tamoxifen induction and genotype of the mouse

Mice were maintained in the animal facility at Kyoto University, and 8-week-old animals were used in this study. Tamoxifen (25 mg/kg in corn oil) was injected intraperitoneally on days -5, - 3, and -1. Recombination of the *Trp53*^*R172H*^ allele was confirmed by PCR analysis of genomic DNA from blood cells of tamoxifen-treated mice. Genotyping of wild-type (290 bp) and rLSL (330 bp) alleles was performed using the primers: Forward 5-AGCCTGCCTAGCTTCCTCAGG-3′ and Reverse 5-CTTGGAGACATAGCCACACTG-3′.

### Immunization by SRBCs and analysis of germinal center reaction by flow cytometer

1x10^8^ SRBCs (Wako, Japan) were injected intraperitoneally (IP) on day 0, followed by a second IP with 1x10^9^ SRBCs. On days 10-12, mice were anesthetized, spleens were collected, then homogenized and filtered through a 40 μm cell strainer to obtain a splenic cell suspension. After red blood cell lysis (420301, Biolegend), the splenocytes were incubated with the antibodies listed in Supplementary Table 1 for 20 minutes at 4 °C in the dark. After washing, the cell suspension was added with 7-AAD (559923, BD, Japan) for 10 minutes before cell sorting by FACS Aria III (BD, Japan).

**Supplementary Table 1:** Antibodies used in FACS.

| Antigens | Product Catalogue number | Titer |
| --- | --- | --- |
| CD45R/B220 | 103257, Biolegend | 1:100 |
| CD185(CXCR5) | 145511, BD, Japan | 1:100 |
| GL7 | 144607, Biolegend | 1:100 |
| CXCR4 | 146507, Biolegend | 1:100 |
| CD83 | 121517, Biolegend | 1:100 |
| CD95(Fas) | 152605, BD Japan | 1:100 |

